# Causal evidence for increased theta and gamma phase consistency in a parieto-frontal network during the maintenance of visual attention

**DOI:** 10.1101/2024.03.17.585395

**Authors:** Claire Bradley, Emily McCann, Abbey S Nydam, Paul Dux, Jason B Mattingley

## Abstract

Endogenous visuo-spatial attention is under the control of a fronto-parietal network of brain regions. One key node in this network, the intra-parietal sulcus (IPS), plays a crucial role in maintaining endogenous attention, but little is known about its ongoing physiology and network dynamics during different attentional states. Here, we investigated the reactivity of the left IPS in response to brain stimulation under different states of selective attention. We recorded electroencephalography (EEG) in response to single pulses of transcranial magnetic stimulation (TMS) of the IPS, while participants (N=44) viewed bilateral random-dot motion displays. Individual MRI-guided TMS pulses targeted the left IPS, while the left primary somatosensory cortex (S1) served as an active control site. In separate blocks of trials, participants were cued to attend covertly to the motion display in one hemifield (left or right) and to report brief coherent motion targets. The perceptual load of the task was manipulated by varying the degree of motion coherence of the targets. Excitability, variability and information content of the neural responses to TMS were assessed by analysing TMS-evoked potential (TEP) amplitude and inter-trial phase clustering (ITPC), and by performing multivariate decoding of attentional state. Results revealed that a left posterior region displayed reduced variability in the phase of theta and gamma oscillations following TMS of the IPS, but not of S1, when attention was directed contralaterally, rather than ipsilaterally to the stimulation site. Under these same conditions, a right frontal cluster also displayed reduced theta variability and increased amplitude of TEPs after TMS of the IPS but not S1. Reliable decoding of attentional state was achieved after TMS pulses of both S1 and IPS. Taken together, our findings suggest that endogenous control of visuo-spatial attention leads to changes in the intrinsic oscillatory properties of the IPS and its associated fronto-parietal network.

## 1. Introduction

Selective attention is the crucial cognitive function that allows adaptive filtering and prioritisation of sensory information based on current goals and stimulus salience (Pashler, 1998). Over the past 30 years, considerable progress has been made in understanding the neural basis of attention, thanks to neuroimaging and electrophysiological investigations (Corbetta & Shulman, 2002; Fiebelkorn & Kastner, 2020; Speed et al., 2020). While most of this research has been correlational in nature, recent use of non-invasive brain stimulation techniques, such as transcranial magnetic stimulation (TMS) (Polanía et al., 2018), has begun to provide causal evidence for the role of fronto-parietal brain areas in regulating attention. However, most of this work has used TMS to interfere with normal brain activity required for successful attentional deployment, which precludes the characterisation of ongoing network dynamics. Here, we combined the strengths of EEG and brain stimulation by recording brain activity in response to single pulses of TMS, to probe the reactivity of a key node in the attention network under different states of selective attention.

According to one influential model, visuo-spatial attention is regulated through the interaction of two complementary networks (Corbetta et al., 2008; Corbetta & Shulman, 2002; Tosoni et al., 2023). The dorsal attention network (DAN) comprises bilateral frontal eye fields (FEF), superior parietal lobules (SPL) and intra-parietal sulci (IPS), whereas the ventral attention network (VAN) consists of the temporoparietal junction and the ventral frontal cortex. Neuroimaging studies have shown that when participants maintain their attention at a given location in space, the DAN is activated, whereas VAN activity is typically reduced (Tosoni et al., 2023), leading to the hypothesis that the DAN is responsible for the allocation and maintenance of endogenous spatial attention. Short bursts of repetitive transcranial magnetic stimulation (rTMS) can be used to temporarily impair neural processing and to establish causal involvement of a given brain area in behaviour. Using this method, key nodes of the DAN - the FEF and ventral IPS – have been shown to be necessary to the maintenance of visuo-spatial attention at a peripheral location, and to the establishment of preparatory oscillatory activity in visual cortices (Capotosto et al., 2009, 2012). In contrast, the SPL has been shown to play a crucial role in shifting attention (Capotosto et al., 2012, 2015). Although the key regions for maintaining endogenous visuo-spatial attention have been identified, much remains to be discovered about how activity within these areas is coordinated.

Repetitive transcranial magnetic stimulation has been instrumental in delineating the role of brain region dynamics and rhythms in visual attention (Capotosto et al., 2009, 2012, 2015; Leitão et al., 2015; Marshall et al., 2015; Riddle et al., 2019; Romei et al., 2012; Wang et al., 2020). However, most studies have relied on disruption or entrainment of brain activity to reveal these effects. This approach makes it hard to investigate the normal physiology of the stimulated area and its associated network during task engagement. Another way to dynamically probe the brain using TMS is to use it not as a means to disrupt ongoing processing, but rather as a means to provide a standard input into the network and to measure the resulting output with concurrent neuroimaging (Bergmann et al., 2016). At rest, this technique has revealed that different cortical areas have distinct resonant properties (Rosanova et al., 2009) and connectivity patterns (Ozdemir et al., 2020), and that an individual’s EEG responses to TMS can be used to establish a ‘cortical fingerprint’ (Ozdemir et al., 2021). Applying this idea to the investigation of cognitive states, it should be possible to probe key brain areas (e.g., using a single pulse of TMS) when the network is in an active state (e.g., attending to a cued location in space), and to measure the resulting network activity when the active state is manipulated (e.g., when the participant is made to attend a different location in space; see Bradley et al., 2022).

For example, in a TMS-fMRI study, Heinen and colleagues (Heinen et al., 2014) presented participants with visual stimuli consisting of faces overlapping with fields of moving dots. When participants were asked to attend to the dots, a TMS probe to the FEF (a key node of the DAN) resulted in greater activity in a motion-sensitive sensory region than in the fusiform face area (FFA), a region specialising in face processing. The opposite pattern of brain activity arose when participants were asked to attend instead to the faces and ignore the moving dots, demonstrating that FEF connectivity varies with the attended feature. While a few other studies have uncovered attention-dependent effective connectivity (Blankenburg et al., 2010; Morishima et al., 2009) and changes in oscillatory activity (Herring et al., 2015; Okazaki et al., 2020), none has provided an in-depth investigation of the effects of attention on the physiology of a node within the DAN following single-pulse TMS. Moreover, TMS-EEG and TMS-fMRI can be combined with machine-learning to assess information content stored in the network being stimulated. In a recent study, Rose and colleagues (Rose et al., 2016) had participants hold two items from separate categories (words, faces or motion signals) in working memory. One item was relevant and ‘attended’ in working memory while the other item was temporarily ‘unattended’ but relevant at a later stage. Following a TMS pulse, the unattended memory item could be transiently decoded from EEG activity, suggesting that TMS pulses can uncover latent category representations coded in patterns of evoked brain activity. To the best of our knowledge, no study has investigated whether TMS pulses can similarly uncover a participant’s latent attentional state.

Here, we asked whether neural activity elicited by parietal TMS pulses varies with visuo-spatial attentional states. We varied spatial attention and attentional load while stimulating a key node of the dorsal attentional system – the left IPS (Silver & Kastner, 2009) – and measuring whole-brain activity with EEG. The IPS was chosen as the stimulation site because it has repeatedly been implicated in maintaining spatial attention at a cued location (Corbetta & Shulman, 2002; Kelley et al., 2008; Sheremata et al., 2018; Szczepanski et al., 2010). The left posterior IPS, in particular, shows stronger contralaterally biased activity relative to the right IPS (Jeong & Xu, 2016; Sheremata et al., 2010, 2018). In our study, we had participants covertly monitor one of two peripheral locations (left or right) for rare events (coherent motion signals in dynamic random-dot displays), thus engaging local activity in support of maintaining visuo-spatial attention. Since activity in the IPS has been shown to be modulated by perceptual load (Xu et al., 2014), we also manipulated load by asking participants to detect stronger or weaker coherent-motion events. TMS pulses were delivered at random times relative to the motion events, resulting in the vast majority of pulses being delivered during periods of random – rather than coherent – motion. This design allowed the sensory aspects of the visual displays to be held constant while varying only the attentional state of the participant (i.e., attend left versus right, and low-load versus high-load). We chose the left primary somatosensory cortex (S1) as an active control site. S1 lies within the parietal lobe, in close proximity to the IPS, so it is well matched to the IPS site in terms of auditory artefacts and muscle activation caused by TMS, while having no known involvement in the allocation of visuospatial attention.

Our overarching aim was to investigate how ongoing attentional state impacts stimulation-induced neural activity. To do so, we focused specifically on measures of electrical brain activity, recorded with EEG in response to the TMS pulse. Imaging studies have shown that attention affects the excitability of visual regions (Corbetta & Shulman, 2002; Poghosyan & Ioannides, 2008) and the variability of neural signals (Arazi et al., 2019). We therefore used TMS-evoked potential (TEP) amplitude as a putative marker of local and remote excitability, and consistency in the phase of oscillatory signals across trials (inter-trial phase clustering, ITPC) as a measure of variability. In addition, we employed multivariate decoding methods to classify attention condition following a TMS pulse, as a measure of global information content (Rose et al., 2016). If current attentional state has no impact on TMS-generated brain activity, then there should be no difference between experimental conditions following TMS over the left IPS, because the visual displays and stimulation intensity were held constant throughout the task. By contrast, if participants’ current attentional state, as defined by spatial location and perceptual load, modulates TMS-evoked brain activity, there should be reliable differences in one or more of these key activity measures across experimental conditions.

## 2. Subjects and methods

### 2.1. Participants and sample size

Forty-six participants (mean age ± SD: 24 ± 3 years; 29 women) took part in the study, which involved a 2-hour session of TMS-EEG and a structural MRI scan used for target localization. Participants received monetary compensation at a rate of $20 per hour for their time, which averaged 3 hours per participant. All participants gave written informed consent prior to taking part and the study was approved by The University of Queensland Human Research Ethics Committee. Participants were screened for contraindications to transcranial magnetic stimulation (TMS), which included a family history of epilepsy, consumption of neuroactive drugs, or a history of neurosurgery or brain injury. All participants completed a standard safety screening questionnaire for TMS (Keel et al., 2001) and for general contraindications to magnetic resonance imaging (MRI). All participants were right-handed, as assessed by the Oldfield handedness questionnaire, and had normal or corrected-to-normal vision. Two participants were excluded prior to analysis: one withdrew from the study before completing both sessions, and another had missing experimental data. This left a final sample size of N=44, as required by an a priori power analysis. The sample size of 44 was chosen to provide 90% power to detect a typical medium effect size of 0.5 when using a paired two-tailed t-test assessed at the standard alpha error probability of 0.05 (see the preregistration on the Open Science Framework https://osf.io/uk86r).

### 2.2. Overview of experimental procedure

Participants attended two experimental sessions. The first was dedicated to screening and structural MRI scanning. The second was dedicated to TMS-EEG. This session involved fitting of the electroencephalography (EEG) cap, TMS neuro-navigation setup, brief training on the behavioural task, followed by motor threshold measurement, TMS-hotspot mapping (of motor, posterior parietal and primary somatosensory sites) and eye-tracker calibration. Participants then performed the motion discrimination task, involving two levels of difficulty (‘Easy’ and ‘Hard’) and two visual spatial attention locations (‘Left’ and ‘Right’) (total duration ∼80 min) (**Fig.1A**). Behaviour (accuracy, reaction time) and EEG activity were recorded throughout. Concurrently to the visual task, but uncorrelated with presentation of visual stimuli, single pulses of TMS were delivered to the posterior parietal cortex (SPL) or to the primary somatosensory cortex (S1), to elicit TMS-evoked potentials (TMS-EPs). The target regions were individually localised as detailed below.

**Figure 1.**
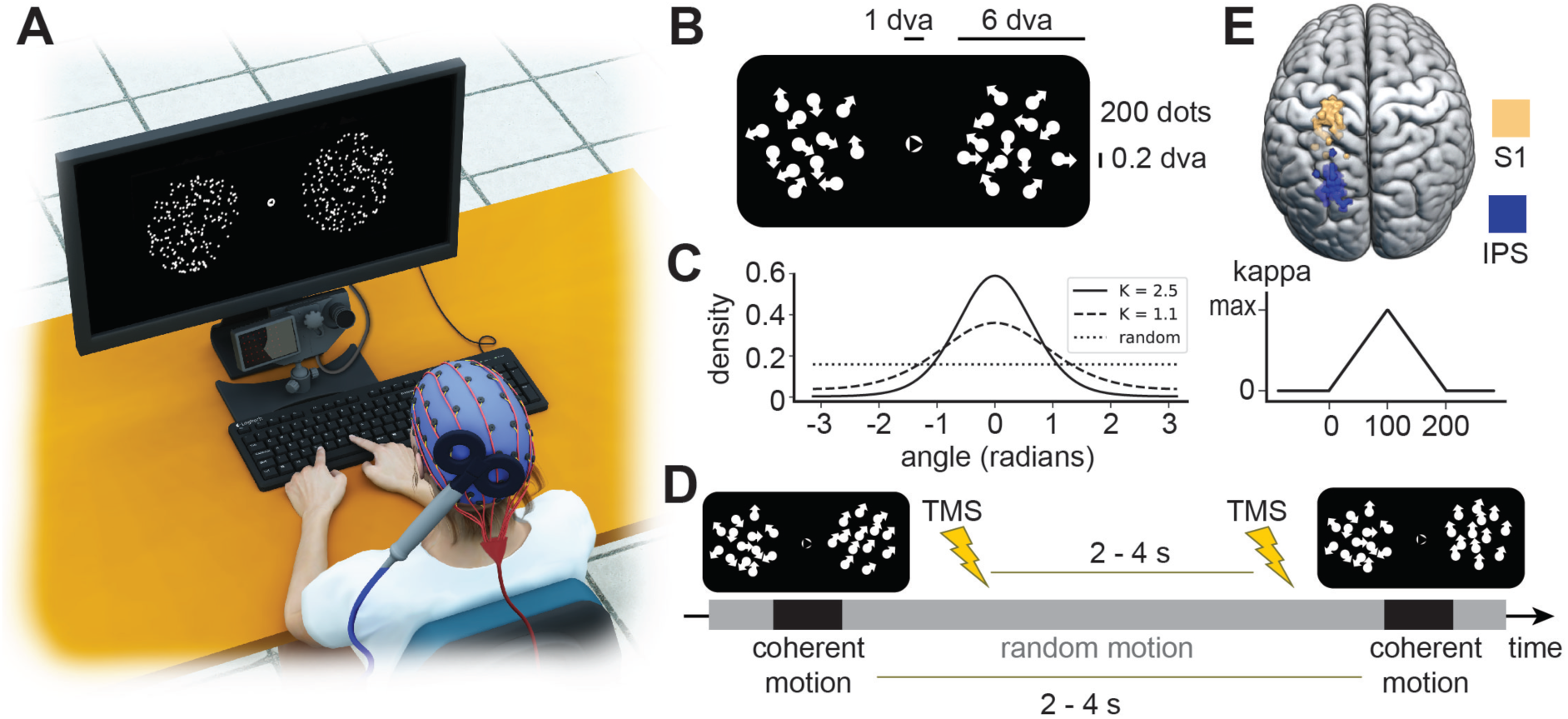
Experimental setup and stimuli. **A)** Participants received transcranial magnetic stimulation (TMS) over the left parietal cortex (black coil), while their brain activity was continuously recorded with a 64-channel electroencephalography system (blue cap). **B)** Stimuli consisted of two patches of moving white dots on a black background, displayed on either side of a white fixation circle. Within the fixation circle, a small black triangle pointed left or right, indicating the side to be covertly monitored for targets with 100% validity. **C)** The direction of the moving dots was taken from one of three von Mises distributions, depending on the condition (*left inset*): a flat distribution resulting in random dot motion, a broad distribution resulting in a weak coherent motion signal (‘Difficult’ condition) or a narrow distribution resulting in a strong coherent motion signal (‘Easy’ condition). To create coherent motion events (*right inset*), the parameter defining distribution width was gradually ramped up and down over 200 ms, giving rise to a brief directed motion signal embedded in random motion. **D)** Coherent motion events occurred randomly every 2 to 4 seconds. Independently of these motion events, single TMS pulses were delivered every 2 to 4 seconds, resulting in 93.8% of TMS pulses being delivered during times of random motion. **E)** TMS pulses targeted one of two regions: the IPS region of the posterior parietal cortex (IPS, blue dots) or the primary somatosensory cortex (S1, yellow dots). Each dot represents the location of stimulation for a single participant, which was individually defined using the participant’s own anatomical MRI scan. N=44 participants.

### 2.3. Eyetracking-controlled covert visual spatial attention task

Participants sat in a chair with their head stabilised by a chin rest and eyes 53 cm from an LCD monitor (ASUS VG248QE; 1920 x 1080 pixels). They were instructed to covertly attend to the cued side while maintaining central fixation, and to press a keyboard key when they detected vertical motion. To avoid a bias in hemispheric EEG activity linked to preparatory motor response activity, participants were requested to respond with the index finger of one hand to vertical upward motion and with the index finger of the other hand for vertical downward motion; the remaining four motion directions were not paired to any response. Reaction time and accuracy were recorded. Participants underwent a short period of training at the beginning of the experiment; they received feedback on their performance at the end of each block and additional verbal encouragement every 8 blocks. To ensure that attention was allocated covertly, eye position was tracked throughout the experiment and monitored online with an EyeLink CL remote-infrared eyetracker (calibration every 8 blocks). Deviations from fixation prompted a reminder to the participant to maintain central fixation.

The visual display consisted of two random dot kinematograms (RDKs) each containing 200 white dots on a black background (6 degrees of visual angle (dva); each dot 0.2 dva), one on each side of a central fixation circle (field eccentricity 10 dva, central circle: 1 dva) (**Fig.1B**). The central fixation circle contained a triangle pointing either left or right, cueing one of the RDKs. Dots moved randomly throughout the task but could transiently move coherently in one of six possible directions (45°, 90°, 135°, 180°, 225°, 270°, 315°). Dot coherence was either high or low, as determined from the width of the distribution from which each dot angle was derived (**Fig.1C**). Briefly, at every frame, dot position was adjusted by a distance and a given angle, which was taken from a von Mises distribution. The von Mises distribution is well suited for continuously varying circular variables, such as angles, and is characterised by a mean (mu) and a width at half maximum (kappa). Here, mu was one of the six angles described above, and the maximum kappa value was 1.1 or 2.5 (low or high coherence, respectively).

The visual task was structured as follows: there were 32 experimental blocks (block length: 25 coherent motion events), which could be from one of 4 pseudo-randomised and counter-balanced experimental conditions: ‘Attend Left, Easy, ‘Attend Right, Easy’, ‘Attend Left, Difficult’ and ‘Attend Right, Difficult’. Attention side and difficulty were announced at the beginning of a block and the fixation arrow served as a reminder for attention side throughout the block. Within a block, both RDKs displayed random motion continuously. On the cued (attention) side, transient events of coherent motion (kappa ramp-up over 100ms and ramp-down over 100 ms) occurred every 2 to 4 seconds (time-steps of 100 ms). There were no coherent motion events on the non-cued side. There was a 20s pause at the end of each block for providing feedback and rest throughout the task. Every 8 blocks, a longer self-paced break was taken, followed by eye-tracker re-calibration.

### 2.4. MRI-guided localisation of TMS targets

Two brain regions were targeted: the left IPS, as the active region of interest to test state-dependency of TMS effects during attentional engagement, and the left primary somatosensory cortex (S1) as a control region. Both IPS and S1 targets were individually localized using the participants’ MRI scans to account for inter-individual variability. T1-weighted magnetization-prepared rapid gradient echo (MPRAGE) images were acquired on a 3T Siemens scanner (TR = 1.9 s, TI = 0.9 s, flip angle = 9°, 3D sequence, 192 sagittal slices, slice thickness = 1mm, in plane voxel size = 1 x 1 x 1 mm, FOV = 256 x 256mm) and segmented in the Visor2 neuro-navigation system (ANT Neuro, The Netherlands) to produce a reconstruction of each participant’s brain, skull and scalp. Middle left IPS was identified on the basis of local sulcal patterns (specifically, the intersection between the main IPS sulcus and a medially pointing sulcus, at the intermediate position between the post-central sulcus and the parieto-occipital fissure, slightly more caudal than IPSa; see Morris et al., 2007); its location (upper bank of sulcal grey matter, on the side inferior and medial to junction) was marked for later stimulation with TMS (**Fig.1E**). Left S1 was defined as the point on the post-central gyrus at about the same lateral location as the left IPS hotspot. Average Montreal Neurological Institute coordinates across participants (+/- SD) were x = -24 +/- 5, y = -78 +/- 6, z = 55 +/- 8 for left IPS, and x = -25 +/- 5, y = -41 +/- 9, z = 79 +/- 5 for left S1.

### 2.5. Neuro-navigation guided TMS

To account for inter-individual differences in cortical excitability and neuroanatomy, TMS intensity for both target and control sites was derived from individual motor cortical threshold, adjusted for the coil-to-cortex distance (Stokes et al., 2005), as described below.

#### 2.5.1. Motor hotspot and threshold measurement

Biphasic TMS pulses were applied through a figure-of-eight coil (Magstim, wing diameter: 70 mm) connected to a Magstim Rapid^2^ magnetic stimulator (Magstim, UK). The coil was held tangentially to the skull with the handle pointing backward and laterally at an angle of 45° to the sagittal plane, at the optimal scalp site to evoke an MEP in the relaxed *abductor pollicis brevis* (APB) muscle of the right hand, as assessed by surface electromyographic (EMG) recordings (for details, see Bradley et al., 2022). Resting motor threshold was defined as the intensity eliciting MEPs above 50 µV peak-to-peak amplitude in at least 10 out of 20 consecutive trials (Rossini et al., 2015). Mean threshold ± SD was 69% ± 11% of maximum stimulator output.

#### 2.5.2. TMS stimulation parameters

Target locations were determined individually, as described above (see ‘*MRI-guided localisation of TMS targets’*). The TMS coil handle was angled ∼30° from the midline. A neuro-navigation system (Visor2) that tracks the position of the coil relative to the participant’s head ensured consistent placement of the TMS coil throughout the experimental session. Taking into account the trade-off between strength of sensory artefacts and certainty of achieving supra-threshold stimulation of a targeted neural population (both higher for high TMS intensities), stimulation intensity (TargetI) was set at 100% of resting motor threshold (rMT). Intensity was adjusted for coil-to-cortex distance as follows: TargetI = 2.7 × (TargetDepth - MotorDepth) + rMT, where TargetDepth is the distance from TMS coil centre to IPS or S1 marker (in mm); MotorDepth is the distance from coil to cortical motor hotspot and rMT is the resting motor threshold (Stokes et al., 2005). Mean stimulation intensity (+/- SD) was 72% +/- 13% of maximum stimulator output for IPS and 75% +/- 12% for S1; intensity was kept constant throughout the experiment. Single biphasic pulses were delivered randomly every 2 to 4 seconds (bins of 250 ms), while participants performed the visual task, with no temporal relationship with the onsets of coherent-motion events. A total of ∼800 pulses (∼100 pulses per block, ∼200 pulses per experimental condition) were delivered over the course of ∼80 minutes. No participant reported any adverse reaction to TMS.

### 2.6. Electroencephalography (EEG) recordings of brain activity

#### 2.6.1. Recording

Continuous EEG data were recorded with a TMS-compatible 64-channel EEG cap (BrainCap, BrainProducts, Germany; Ag/AgCl electrodes) in accordance with the 10–10 extended international system. All electrodes were referred to the right mastoid and impedance was kept below 5 kΩ using a viscous electrode gel (SignaGel, Parker; skin preparation with NuPrep, Weaver & Co); the ground electrode was incorporated in the cap at the AFz electrode site. BrainRecorder software and BrainAmp MR Plus amplifiers (BrainProducts, Germany) were used to record EEG with a 5000 Hz sampling rate and low-pass filter of 1000 Hz (no high-pass filter, resulting in a DC recording).

#### 2.6.2. TMS-EPs preprocessing

Pre-processing of EEG data containing TMS-linked activity is challenging – among other things due to the presence of artefacts several orders of magnitude larger than brain activity – and requires special processing steps (Hernandez-Pavon et al., 2023; Rogasch et al., 2017). All analyses were performed in Matlab, using EEGLAB (v13.6.5b, Delorme & Makeig, 2004) and the TESA toolbox (Rogasch et al., 2017). Briefly, continuous data were segmented relative to TMS pulse onset (-1500ms to 1500ms) and down-sampled from 5000Hz to 1000Hz. Baseline correction (-1500ms to -10ms) was applied before performing manual trial rejection. Two electrodes (TP9 and TP10) were excluded because of systematic noise; other noisy or flat electrodes were eliminated based on their filtered (1-100Hz band-pass, 50Hz notch) activity during the baseline (-1500ms to 0ms) and later interpolated. Separate ICA decompositions were performed on data from all blocks for the two TMS stimulation sites (i.e., separate ICAs for IPS and S1 targets) because different stimulation artefacts were expected. The first ICA eliminated large TMS-related artefacts only, the second eliminated a variety of remaining artefacts (i.e., muscle activity, eye blinks, electrode noise, etc.); component classification and rejection was also validated by an experimenter. Data were filtered (1-100Hz band-pass, 50Hz notch) before the second ICA. Electrodes were interpolated and signals re-referenced to the average after the second ICA. Data from different blocks were averaged to yield a TEP for each experimental condition, as well as grand-average TEPs for IPS and S1 target conditions, and partial grand-averages for attention conditions (Attend Left, Attend Right, High coherence, Low coherence).

### 2.7. Time-locked TMS-EPs analysis

#### 2.7.1. Peak analysis

Peaks were detected within a time-window of variable size based on peak latency (10 ms for peaks under 40 ms, 20 ms for peaks between 40 and 140ms, and 80ms for peaks beyond 140ms), centered on a local maximum of global mean field potential (GMFP), as described in details in the Results (see below).

#### 2.7.2. Cluster-based permutation testing

Cluster-based permutation testing (Oostenveld et al., 2011) was performed in the MNE-Python toolbox (Gramfort et al., 2013; Larson et al., 2024) (sample statistic: dependent samples t-test; 1000 permutations, cluster-forming alpha level: 0.001, two-tailed test alpha level: 0.05) on whole scalp data to compare categories for Attention Side (‘Left’ and ‘Right’) and Difficulty (‘Hard’ and ‘Easy’) separately, following IPS and S1 stimulation.

### 2.8. Inter-trial phase clustering and power analyses

EEG signals can be analysed not only in the time domain, as described above, but also in the frequency domain, where signals are decomposed into a combination of oscillatory activities at different frequencies. We investigated variability in the frequency domain by looking at inter-trial phase clustering (ITPC, Cohen, 2014). ITPC measures how consistent the phase of the oscillatory activity is across trials. These measures are best interpreted together with power values, because changes in power may confound ITPC evaluation (van Diepen & Mazaheri, 2018). We therefore also investigated oscillatory power across experimental conditions.

All analyses were performed in Python with the MNE-Python toolbox. Because ITPC measures are generally sensitive to differences in trial numbers, single trial epoch numbers were equalized within each condition (‘Attention Side’, ‘Difficulty’) for both relevant categories and stimulation sites. Individual average power and inter-trial phase clustering values were extracted from single trials using Morlet wavelets centered on frequency bins ranging from 4 Hz to 45 Hz (1Hz steps, cycles per wavelet: frequency/2) and data were decimated by a factor of 3 to reduce memory usage. Power values were baseline corrected with a logratio correction using the -1500 to 0 ms time-interval. Differences between categories in a condition were investigated using cluster-based permutation testing over time-frequency-channel space (sample statistic: dependent samples t-test; 1000 permutations, cluster-forming alpha level: 0.001, two-tailed test alpha level: 0.05) on whole scalp data, separately for IPS and S1 stimulation sites.

### 2.9. Decoding analysis

In order to determine whether TMS pulses uncovered latent attentional state information (Rose et al., 2016), we used a multivariate decoding approach of whole-scalp EEG (Grootswagers et al., 2017). In cognitive neuroscience, multivariate decoding is often used to determine whether a given pattern of brain activity is associated with a given perceptual, cognitive or motor state versus another. It takes into account information from many sources, rather than from a single measure, and can thus be more sensitive to distributed patterns that are not well captured by univariate analyses. In the context of TMS-EEG, decoding has recently been applied to the prediction of motor cortex excitability states, as defined by motor evoked potentials (Hussain & Quentin, 2022; Metsomaa et al., 2021). Here, we sought to characterise participants’ attentional state, as defined by the location of spatial attention and perceptual load, from TMS-evoked potentials. Decoding analyses were performed in MATLAB, using the CoSMoMVPA toolbox (Oosterhof et al., 2016). For a given condition (e.g. ‘Attention Side’), EEG data were downsampled to 200 Hz to speed up computation time, trials were labelled as belonging to one of two category (e.g., ‘Right’ or ‘Left’), and were averaged 4 by 4 to increase signal-to-noise ratio (Grootswagers et al., 2017). A linear discriminant analysis (LDA) classifier was applied to each time-point separately, using a four-fold cross-validation scheme with no repeats. Briefly, this involved training the classifier on three-quarters of the trials and testing its performance on the remaining quarter of trials, and repeating this until all trials had served as test trials. Average classification accuracy over four folds for each participant was then smoothed (5 time-points moving average) and submitted to Bayesian statistical analysis. The Bayesian equivalent of a t-test at each time-point was used to determine whether there was evidence in favour of, or against, the alternative hypothesis that decoding was above chance (Matlab wrapper for package BayesFactor, (Morey & Rouder, 2018). A JZS prior with a scale factor of 0.707 was used, since it is a default in the BayesFactor package that makes minimal assumptions about effect sizes (Teichmann et al., 2022). Following published recommendations (Teichmann et al., 2022), the null hypothesis of nil effect size (d=0) was compared to the alternative hypothesis interval of effect sizes [0.5 to Inf[, equivalent to a one-sided t-test. that medium to large positive effect sizes were expected, while small effect sizes were deemed not relevant. This was done separately for the left IPS and S1 stimulation sites. Bayes Factors (BF) were interpreted following Lee & Wagenmakers, 2014, with BF smaller than 1/3 providing moderate evidence for the null hypothesis and BF larger than 10 providing strong evidence in favour of the alternative hypothesis.

## 3. Results

### 3.1. Behaviour

First, we analysed behavioural variables to verify that the task difficulty manipulation was effective, and to ensure that there were no differences in performance between the attended RDKs on the left and right sides. Because our aim was not to use TMS to probe relevant brain networks, rather than to disrupt behaviour, we verified that there were no differential effects of TMS on accuracy and reaction times across conditions. Accuracy was defined as the proportion of correctly detected targets relative to the number of target trials overall. A correct trial was defined as a trial in which the participant pressed the appropriate key for an upward or downward motion event within a 2s window following that motion event. Pressing the wrong key, responding after 2s, or pressing the key in response to non-target motion events were all labelled as incorrect responses and were excluded. Reaction time (RT) was defined for correct trials only. Accuracy and reaction time were subjected to a 2 x 2 x 2 repeated-measures analysis of variance (ANOVA), with factors of Attention Side (‘Left’ and ‘Right’), Difficulty (‘Hard’ and ‘Easy’) and TMS site (‘IPS’ and ‘S1’).

As expected, participants’ accuracy was lower for low-coherence (hard) targets (44% hits) than high-coherence (easy) targets (81% hits), (F(1,43)=430.4, p<0.0001). There were no significant main effects of Attention side or TMS site, and no interaction between these factors (all F values < 2.3). Also as expected, participants’ RTs were slower for low-coherence (hard) targets (mean +/- SEM: 917 +/- 22ms) than high-coherence (easy) targets (840 +/- 18ms), (F(1,43)=59.15, p=<0.0001). Again, there were no significant effects of Attention side or TMS site for RTs, and no interaction between these factors (all F values < 1.8). Taken together, these results confirm that the task’s difficulty manipulation was effective, and that TMS pulses did not affect performance differentially between the left and right hemifields.

### 3.2. Grand-average TMS-evoked potentials (TEPs)

We next investigated the effect of attentional state, as manipulated by the blocked spatial attention and motion-coherence manipulations, on neural responses to focal TMS pulses over IPS and S1. To do so, we averaged electrical brain activity following a TMS pulse over hundreds of trials. The resulting TMS-evoked potentials (TEPs) comprise a series of peaks and troughs which are specific to the site of stimulation (**Fig.2**) and which have been shown to reflect activation of the stimulated site and interconnected regions (refs). Differences in TEPs between conditions were assessed following two approaches: a classical peak-based analysis, and mass-univariate cluster-based permutation testing.

**Figure 2.**
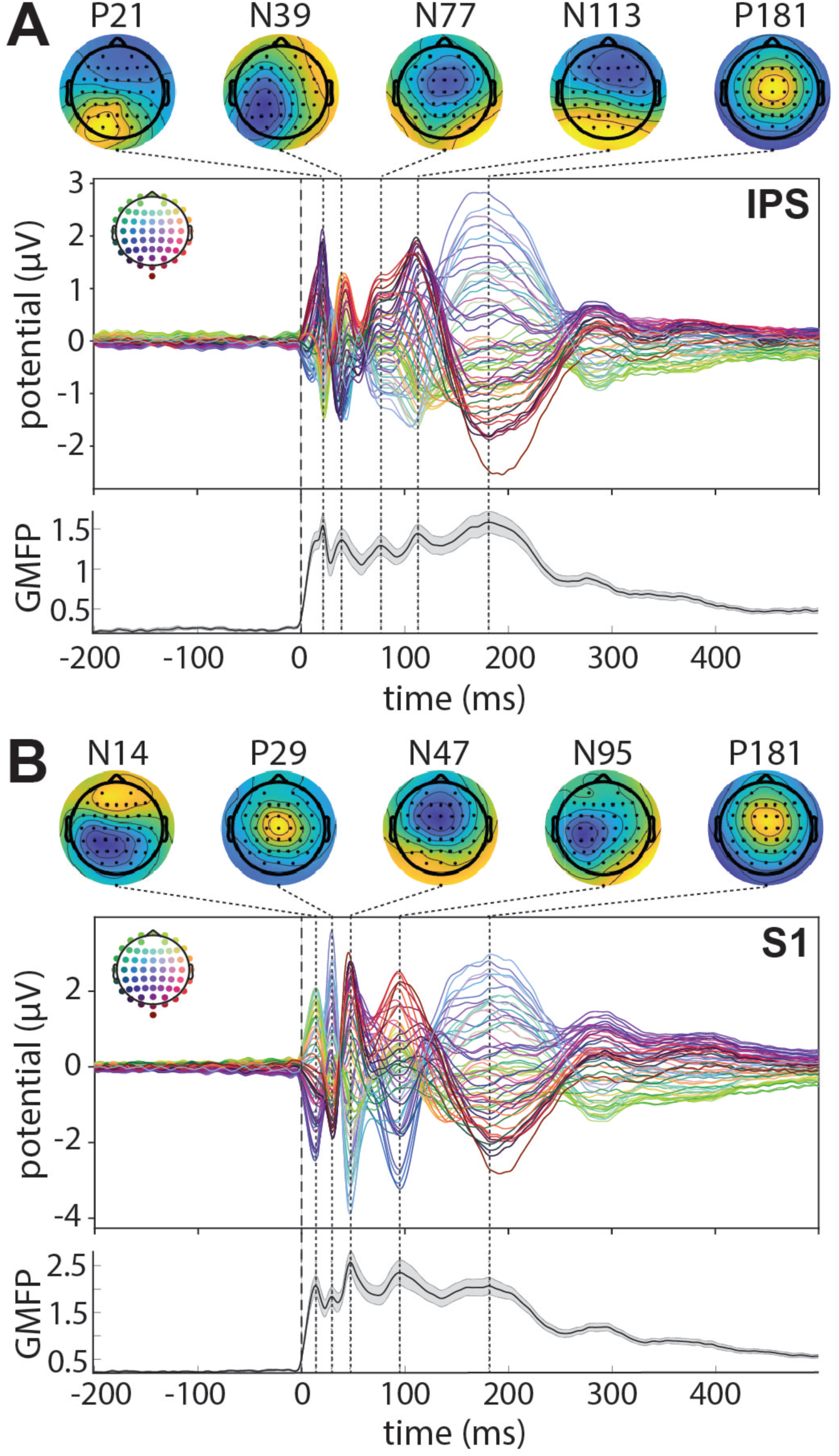
Grand-average TMS-evoked potentials (TEPs). Data shown for TMS delivered over the left intraparietal sulcus (IPS) site **(A)** and the primary somatosensory (S1) control site **(B)**. Voltage over time at each scalp EEG electrode – shown in the *central panel* - is colour-coded according to a spatial distribution shown in the head-map inset, where each coloured dot represents an electrode. Time zero (*vertical dashed line*) is the time of delivery of the TMS pulse. Vertical dotted lines indicate the times of peak global mean field potential (GMFP), shown in the *lower panel* as the average over participants, with standard error of the mean shown in grey shading. Topographic maps at each of the GMFP peaks are shown in the *upper panel*, scaled to the local minimum (blue colour) and maximum (yellow colour). Each black dot represents an electrode and the labels above each map describe the polarity of the peak (N for negative, P for positive) and its latency (in ms). N=44 participants.

### 3.3. Peak amplitude

Five peaks were defined based on local maxima in the Global Mean Field Potential (GMFP), a summary measure reflecting the first derivative of voltage at all electrodes (Skrandies, 1990). GMFP was calculated from the grand-average of all conditions and participants, separately for each TMS site (**Fig.2A, 2B, lower panels**).

A time-window centered around each GMFP maximum was defined, in which to search for an EEG peak (Rogasch et al., 2017). For each local maximum, the electrodes that made up the dominant topographic pattern (negative or positive) (**Fig.2, upper panels**) were selected; voltage from these electrodes was then averaged. Peak amplitude (absolute maximum amplitude within a time-window) was then extracted from this averaged voltage trace. Statistical analysis of peak amplitudes consisted of two-sided, paired t-tests for ‘Attention Side’ (‘Left’ vs ‘Right’) and ‘Difficulty’ (‘Hard’ vs ‘Easy’). Significance level was set to p=0.05, before applying a Bonferroni correction (p=0.0025 for the 5 peaks x 2 conditions x 2 sites explored).

For IPS stimulation, P181 amplitude was larger in the ‘Hard’ (mean +/- SEM: 2.91 +/- 0.27μV) than in the ‘Easy’ condition (2.63 +/- 0.26μV), (t(43)=-3.78, p<0.001). There was no significant effect of difficulty on any other peak (all p>0.05). There was no significant effect of Attention Side on any peak (all p>0.05). For S1 stimulation, P181 amplitude was larger in the ‘Hard’ (mean +/- SEM: 3.19 +/- 0.34μV) than in the ‘Easy’ condition (2.78 +/- 0.31μV), (t(41)=-4.39, p<0.0001, after exclusion of two outliers). There was no significant effect of Difficulty on any other peak after correction for multiple comparisons (all p>0.005). There was also no significant effect of Attention Side on any peaks after correction for multiple comparisons (all p>0.01). These results suggest that task difficulty modulates a ‘late’ TMS-evoked potential (P181) following both IPS and S1 stimulation.

### 3.4. Cluster-based permutation testing

Cluster-based permutation testing (CBPT) is a statistical method that takes into account all EEG data instead of restricting analysis to a few peak windows, while correcting for multiple comparisons (Maris & Oostenveld, 2007). CBPT involves running statistical tests at all time-points and all electrodes and then finding ‘clusters’ of test results above a certain threshold that are adjacent in time and space. The size of these clusters is non-parametrically compared to that of clusters formed from many permutations of the two conditions. The probability attached to this comparison (p-values reported throughout the text) testing of the null hypothesis that the two conditions are exchangeable. It is important to note that a given cluster cannot not be interpreted as ‘the’ significant difference in the data; rather, it is a portion of what contributed to the significant difference between conditions (Sassenhagen & Draschkow, 2019).

We applied a CBPT to voltage data at the whole scalp level in the latency range from 0 to 300 ms post-TMS pulse to investigate the effect of Attention Side and Difficulty on TEPs. **Figure 3** shows the spatial and temporal extent of clusters found when a significant difference between conditions emerged.

**Figure 3.**
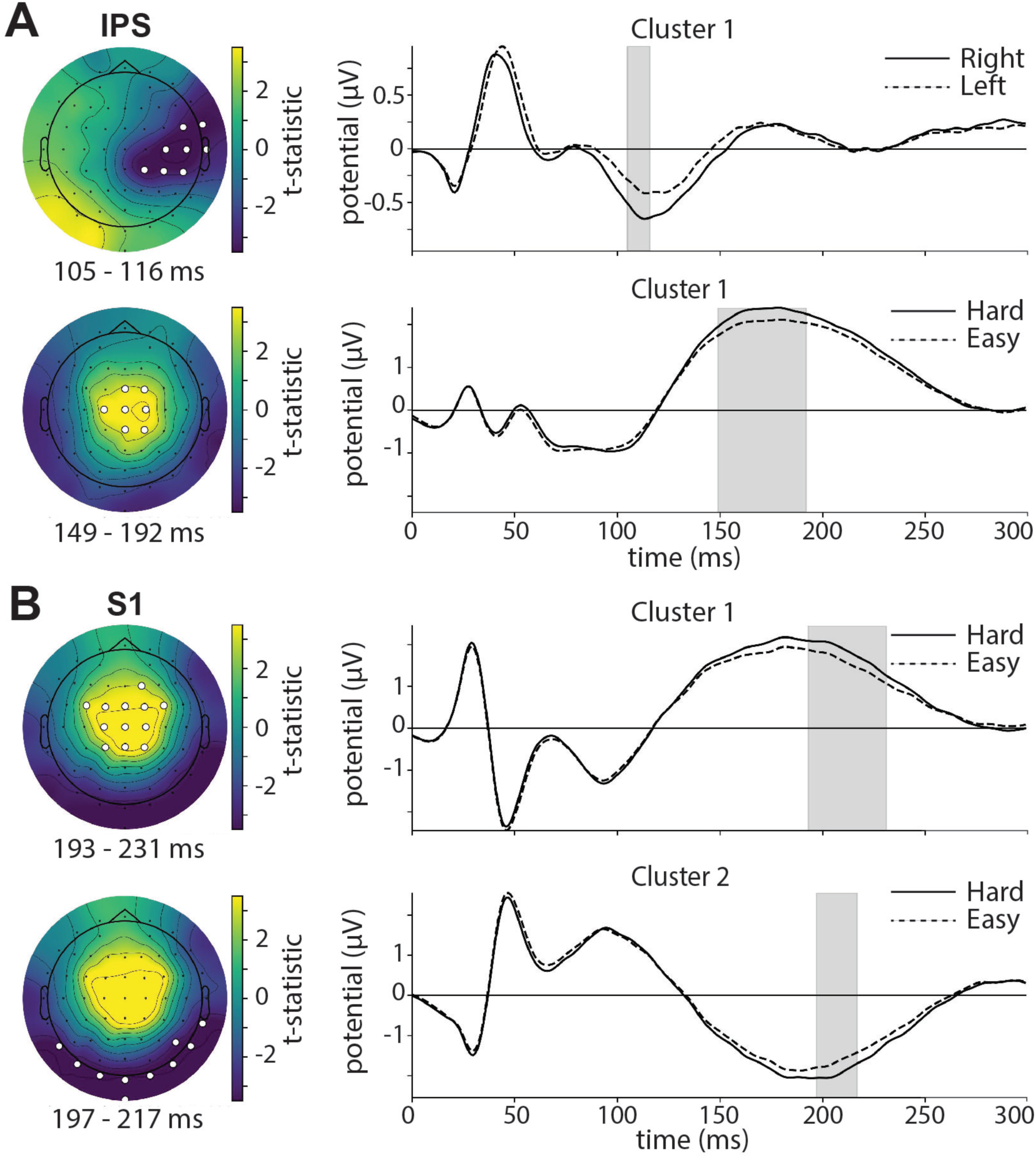
Difference in voltage over time as a function of Attention Side (Left, Right) and Difficulty (Easy, Hard). **A)** TMS delivered over posterior parietal cortex (IPS). **B)** TMS delivered over primary somatosensory cortex (S1). Significant clusters identified by cluster-based permutation testing are shown, in the form of topographic maps (left) and time-domain representations (right). The maps display the t-statistic for the contrast of interest (‘Attend Right – Attend Left’, or ‘Hard – Easy’). Electrodes belonging to a cluster are highlighted in white, and the time range of a cluster is highlighted by a grey shaded rectangle. Each map is the average t-statistic over the time range of its corresponding cluster, and each time-domain trace is the average voltage at the highlighted electrodes. Time zero is the time of delivery of a TMS pulse. N=44 participants.

For IPS stimulation, there was a significant main effect of Attention Side (p<0.05), reflecting a larger negative potential in the ‘Attend Right’ (contralateral to TMS) than in the ‘Attend Left’ (ipsilateral to TMS) condition in a cluster of right fronto-central electrodes around 110ms after TMS (**Fig.3A**, top). There was also a significant main effect of Difficulty (p<0.01), which was driven in part by a larger positive potential for ‘Hard’ than ‘Easy’ trials at central electrodes around 170ms after TMS (**Fig.3A**, bottom). For S1 stimulation, there was no significant main effect of Attention Side, but there was a significant effect of Difficulty (two clusters, p<0.01). There was a larger positive potential for ‘Hard’ than ‘Easy’ trials at central electrodes around 210ms after TMS, and a larger negative potential for ‘Hard’ than ‘Easy’ trials at posterior electrodes around 210ms (**Fig.3B**). These results corroborate the peak analysis in revealing an effect of task difficulty on ‘late’ potentials following both IPS and S1 stimulation. They also show an effect of attention side exclusively following IPS stimulation.

### 3.5. ITPC & Power

While attention can modulate the overall strength of neuronal responses (Corbetta & Shulman, 2002; Poghosyan & Ioannides, 2008), it can also affect the variability of such responses (Arazi et al., 2019). Variability in responses to TMS as a function of attention has rarely been investigated experimentally, despite potentially co-varying with ongoing functional state (Noreika et al., 2020). In order to explore neural variability following TMS across attention conditions, we investigated the alignment of phase across trials (inter-trial phase clustering, ITPC). To this end, we decomposed EEG activity into oscillatory components in the 4-45Hz range and evaluated their phase and power over time. We analysed the effects of Attention Side (**Fig.4**) and Difficulty (**Fig.5**) on phase and power using cluster-based permutation testing at the whole scalp level, over the whole frequency range, in the latency range from 0 to 1000ms post-TMS pulse. Of note, ITPC modulation in the absence of power changes can safely be interpreted as true ITPC modulation, whereas an ITPC modulation together with power modulation in an overlapping time-frequency range should be interpreted with caution (van Diepen and Mazaheri 2018).

**Figure 4.**
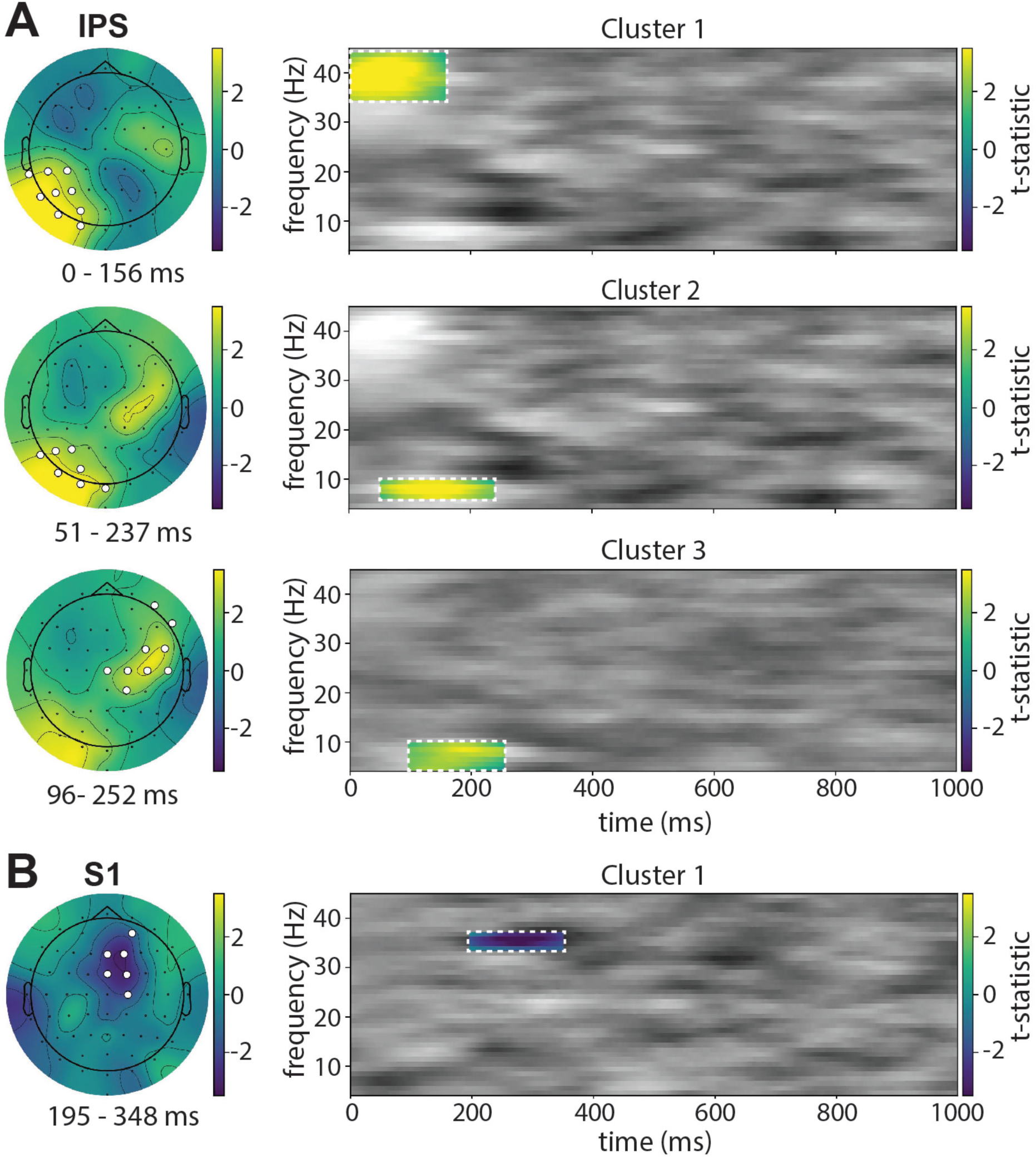
Difference in inter-trial phase consistency (ITPC) as a function of Attention Side. **A)** TMS delivered over the intraparietal sulcus (IPS). **B)** TMS delivered over primary somatosensory cortex (S1). Significant clusters following cluster-based permutation testing are shown, in the form of topographic maps (left) and time-frequency representations (right). Both maps and time-frequency representations display the t-statistic for the contrast ‘Attend Right – Attend Left’. Electrodes belonging to a cluster are highlighted in white, and the frequency and time ranges of a cluster are delineated by white dashed lines (highlighted in colour). Each map is the average t-statistic over the time and frequency range of its corresponding cluster, and each time-frequency map is the average t-statistic at the highlighted electrodes. Time zero is the time of delivery of a TMS pulse. N=44 participants.

**Figure 5.**
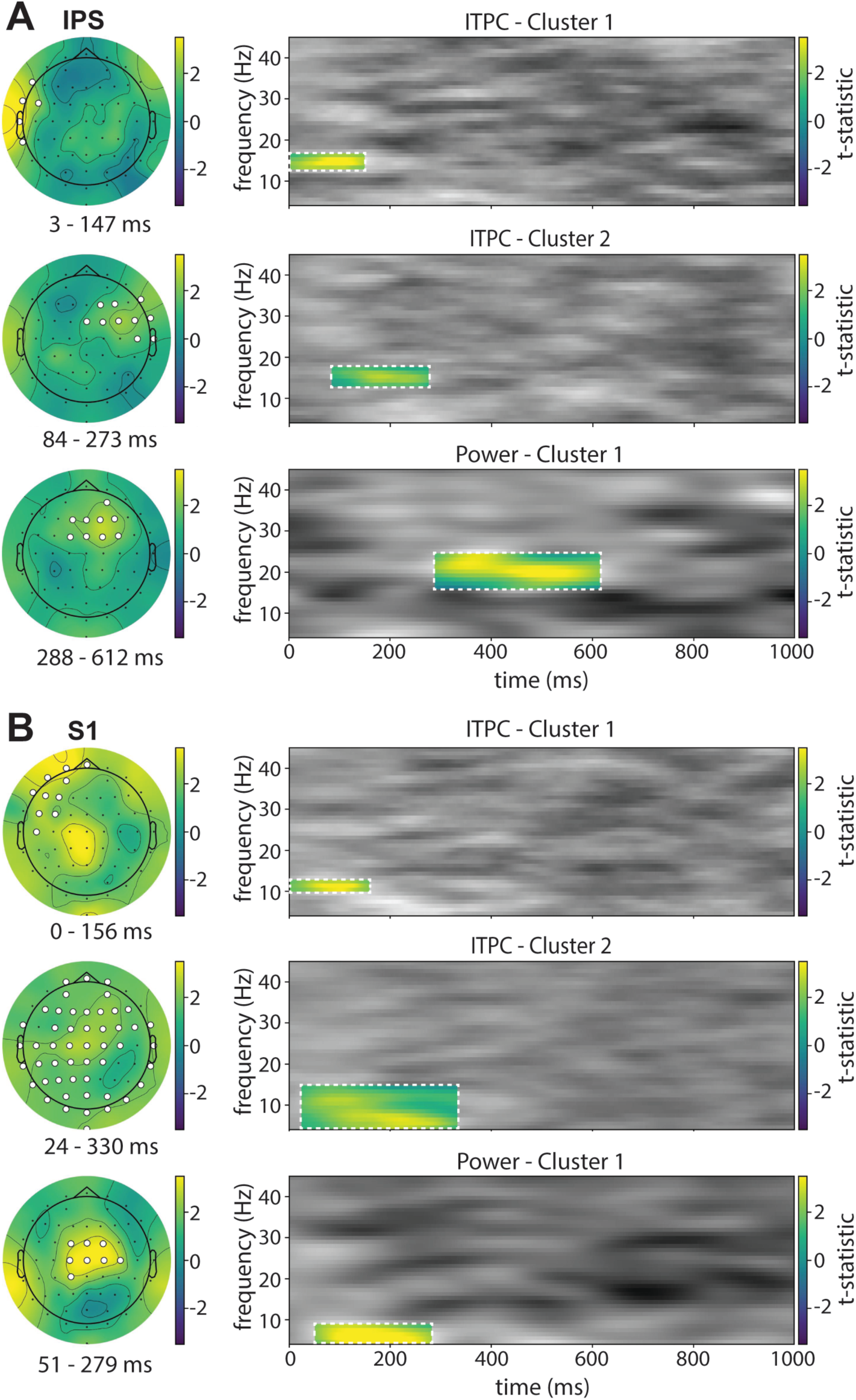
Difference in inter-trial phase consistency (ITPC) and power as a function of task difficulty. **A)** TMS delivered over the intraparietal sulcus (IPS). **B)** TMS delivered over primary somatosensory cortex (S1). Significant clusters following cluster-based permutation testing are shown, in the form of topographic maps (left) and time-frequency representations (right). Both maps and time-frequency representations display the t-statistic for the contrast ‘Hard – Easy’. Electrodes belonging to a cluster are highlighted in white, and the frequency and time ranges of a cluster are delineated by white dashed lines (highlighted in colour). Each map is the average data over time and frequency ranges of its corresponding cluster, and each time-frequency map is the average data at the highlighted electrodes. Time zero is the time of delivery of a TMS pulse. N=44 participants.

For IPS stimulation, there was a significant main effect of Attention Side. Three clusters emerged, showing higher ITPC (i.e., *less variability* in timing across trials) in the ‘Attend Right’ condition (contralateral to TMS) than in the ‘Attend Left’ condition (ipsilateral to TMS) (**Fig.4A**). Cluster 1 (p=0.001) involved left posterior parietal electrodes near the stimulation site, in the gamma range (∼40 Hz), around 0-150ms after TMS. Cluster 2 (p<0.01) involved a similar cluster of electrodes, in the theta range (6-9 Hz), around 50-240ms. Cluster 3 (p=0.01) involved right fronto-central electrodes, in the theta range (4-9 Hz), around 95-250ms. There was no significant main effect of Attention Side when considering power. For S1 stimulation, a cluster-based permutation test of ITPC data in the latency range from 0 to 1000ms revealed an effect of Attention Side. In that time-window, one cluster emerged (p<0.05), which showed lower ITPC (i.e., *more variability* in timing across trials) in the ‘Attend Right’ condition (contralateral to TMS) than in the ‘Attend Left’ condition (ipsilateral to TMS) in a group of fronto-central electrodes, in the low gamma range (∼35 Hz), around 190-350ms after the TMS pulse (**Fig.4B**). There was no main significant effect of Attention Side when considering power.

For the effect of Difficulty, two significant clusters emerged following IPS stimulation, reflecting higher ITPC (i.e., *less variability* in timing across trials) in the ‘Hard’ than in the ‘Easy’ condition (**Fig.5A**). Cluster 1 (p<0.05) involved left temporal electrodes, in the beta range (∼15 Hz), around 0-150ms following TMS. Cluster 2 (p<0.05) involved a right fronto-central set of electrodes, in the beta range (∼15 Hz), around 85-270ms after TMS. These clusters were accompanied by a significant effect of difficulty on power (p<0.01), as driven by a fronto-central cluster at 20 Hz around 290-610ms, where power was higher in the ‘Hard’ than in the ‘Easy’ condition.

For S1 stimulation, there was a significant effect of difficulty involving two clusters, both showing higher ITPC (i.e., *less variability* in timing across trials) in the ‘Hard’ than in the ‘Easy’ condition (**Fig.5B**). Cluster 1 (p<0.05) involved left fronto-temporal electrodes, in the alpha range (10-13 Hz), around 0-150ms after TMS. Cluster 2 (p=0.001) involved a majority of the scalp electrodes, in the theta to alpha range (4-14 Hz), around 20-330ms after TMS. This finding was accompanied by a significant effect of difficulty on power (p=0.01), as driven by a central cluster at 4-8 Hz from 50-280ms, where power was higher in the ‘Hard’ than in the ‘Easy’ condition.

### 3.6. Decoding

In a final step, we applied a multivariate decoding approach to the voltage data in order to investigate whether TMS pulses uncovered latent neural states, as has been shown for working memory tasks (see Rose et al., 2016). The rationale for this analysis was to take voltage information at one time-point across the whole scalp and to use it to train a classifier to distinguish between task states (i.e., between the two levels of Attention Side (Left vs Right) and two levels of Difficulty (Easy vs Hard)). If the classifier is able to correctly label new samples of left-out TMS-evoked neural responses, it means that state-related information has been triggered by the TMS pulses. Evidence in favor of, or against, the classifier performing above chance was evaluated using Bayesian statistics.

**Figure 6** shows decoding accuracy over time for both of the task conditions and TMS target sites. Overall, decoding accuracy was at chance level before TMS pulses, and briefly rose to significantly exceed chance after a TMS pulse, in both conditions and for both TMS target sites.

**Figure 6.**
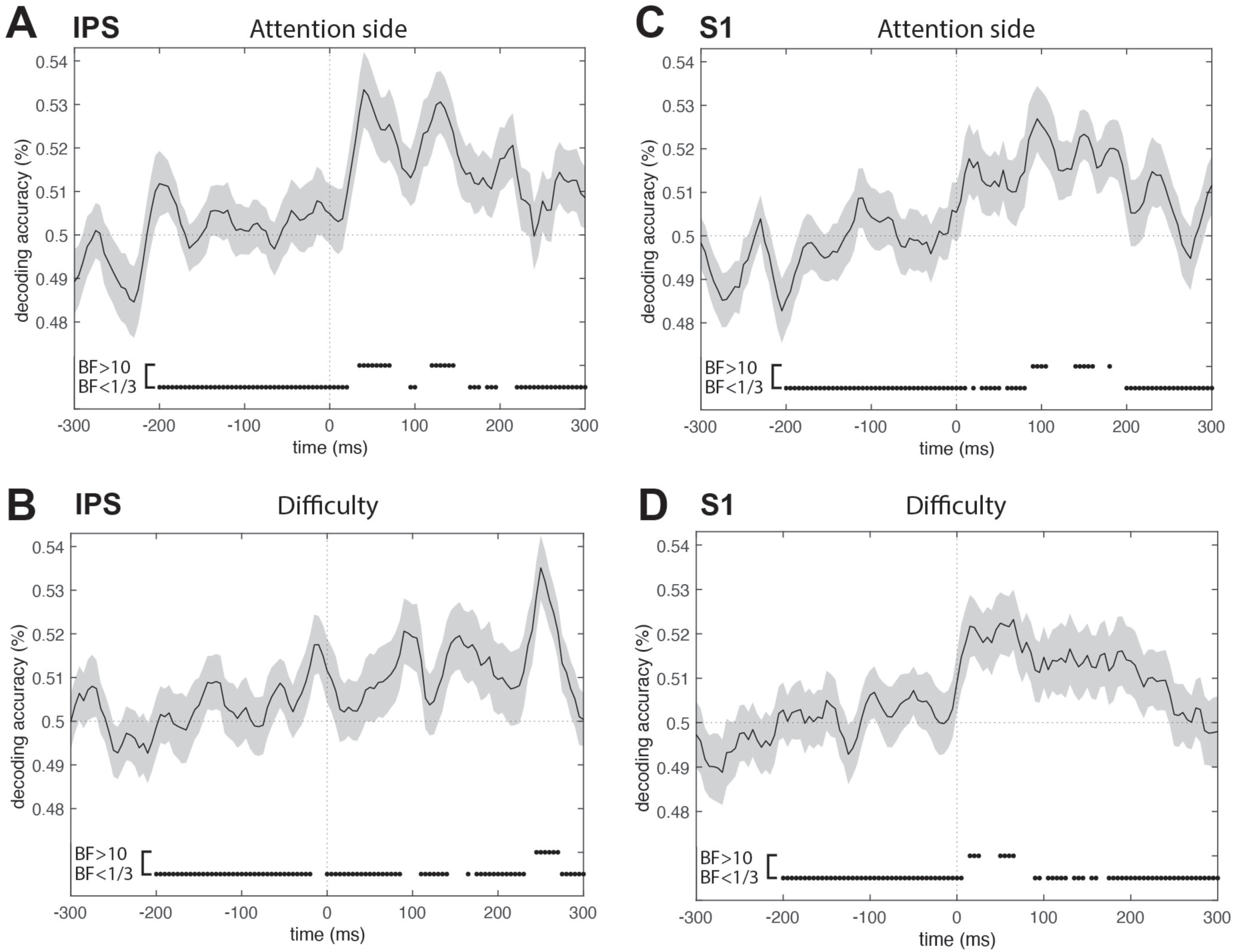
Decoding experimental condition from whole-scalp voltage, following IPS stimulation **(A, B)** and S1 stimulation (C, D). A Linear Discriminant Analysis (LDA) was applied at each time-point to classify whole-scalp voltage data as belonging to the ‘Attend Left’ or ‘Attend Right’ condition **(A, C)** or to the ‘Hard’ or ‘Easy’ condition **(B, D)**. Results are presented as average decoding accuracy over participants, with standard error of the mean as grey shading. The horizontal dotted line denotes chance level; the vertical dotted line denotes the time of TMS pulse delivery. Bayes Factors (BF) in favour of (top, BF>10) or against (bottom, BF<1/3) rejecting the null hypothesis of no above-chance decoding are shown in an *inset* below the main plots. N=44 participants.

For IPS stimulation (left panels), decoding accuracy was at chance level prior to the TMS pulse (Bayes Factors in favor of the null hypothesis of ‘no difference from 50% accuracy’ shown below the accuracy time-course). Following a TMS pulse, there was evidence in favor of the alternative hypothesis (‘accuracy is different from 50%’) at around 50 ms and 130 ms for ‘Attention Side’ and at around 100 ms and 260 ms for ‘Difficulty’. This indicates that TMS pulses delivered over the left IPS transiently uncovered latent information about the spatial attentional state of the participant. For S1 stimulation (right panels), decoding accuracy was similarly at chance level prior to a TMS pulse. Following a TMS pulse, there was evidence in favor of the alternative hypothesis at around 100 to 170 ms for ‘Attention Side’ and around 10 to 60 ms for ‘Difficulty’.

## 4. Discussion

The overarching goal of this study was to investigate the role of attentional state in modulating neural responses to brain stimulation of the left intra-parietal sulcus (IPS), a key node within the dorsal attention network (DAN). Previous research has shown attention-dependent changes in effective connectivity between different nodes in the attentional network and visual areas (Blankenburg et al., 2010; Heinen et al., 2014; Morishima et al., 2009), coupled with changes in oscillatory patterns in visual cortex (Herring et al., 2015; Okazaki et al., 2020). Despite the central role of IPS in visual attention, no research has specifically examined the effects of attention on the excitability, variability and information content of TMS-evoked responses originating from IPS during its engagement in a visual attention task. We tested the specificity of such effects by stimulating an active control site, left S1, which was not involved in the task at hand. A number of results reveal state-dependent effects that are specific to IPS stimulation. A cluster of right frontal electrodes showed larger amplitude TEPs when attention was directed contralaterally to the site of IPS stimulation, as opposed to ipsilaterally. That same cluster, along with electrodes proximal to the stimulation site, also displayed less variability in the phase of theta oscillations for contralateral versus ipsilateral spatial attention. A left posterior cluster also showed reduced gamma variability under these conditions. Such effects were not seen following S1 stimulation. Other measures showed no specific differences between IPS and S1 stimulation. TEP peak amplitude generally did not differ by attentional state, but when it did, a similar increase was found for both S1 and IPS stimulation. Similarly, it was possible to decode attention side and task difficulty, regardless of whether the TMS pulse was delivered to the IPS or S1, albeit with different time-courses for the two areas of stimulation.

A number of design elements allowed us to rule out that the observed differences between conditions might have arisen due to changes other than task-modulated attentional state. First, the TMS pulses were delivered predominantly during periods in which only random motion was present in the displays, rather than during the brief intervals containing coherent motion targets. This ensured that sensory inputs from the two motion patches remained constant at the time of TMS evoked potentials. All that changed across conditions was whether participants attended to one side of space or the other, and whether they anticipated low- or high-coherence target stimuli. Second, attention was always deployed covertly by participants, and eye position was verified using eye-tracking, thereby ruling out any confounding effects of changing eye position across experimental conditions. Third, the location of the stimulating coil was monitored with an infra-red camera and held constant over either the left IPS or left S1 (active control) region. Finally, the experimental conditions were delivered in short blocks that were counterbalanced at the individual level, ruling out any biases that might have arisen from slow, session-wide changes in evoked potentials or effects produced by practice or fatigue.

### Visuo-spatial attention reduces inter-trial variability in a fronto-parietal network

We observed an effect of spatial attention on inter-trial phase clustering (ITPC) that was different for IPS stimulation versus S1 stimulation. Notably, following left IPS stimulation, a cluster of electrodes over the left posterior scalp showed higher gamma and theta ITPC (i.e., less variability) when attention was directed contralaterally rather than ipsilaterally to the stimulated (left) hemisphere. Concurrently, a cluster of electrodes over the right frontal region also showed higher ITPC in the theta range. Although scalp potentials can originate from brain regions not directly underlying the electrodes (Michel & He, 2019), the potentials we observed are broadly consistent with activity in the left posterior parietal, left occipital and right frontal cortices. These findings show that spatial attention is associated with less theta band variability in a fronto-parietal network when attention is directed contralaterally rather than ipsilaterally to the hemisphere of stimulation. This is interesting because there is growing evidence for a theta-rhythmic modulation of behaviour and neural activity that is attributable to attentional sampling (Fiebelkorn & Kastner, 2020). According to this account, the phase of theta activity in a fronto-parietal network is linked to two states, which allow for alternating periods of higher performance and attentional focus, and lower performance coupled with an opportunity to change the focus of attention (Fiebelkorn & Kastner, 2020). Our results suggest that attentional engagement contralateral to the stimulated IPS allowed the TMS pulse to reset oscillation phase in a distributed network more efficiently than when attention was engaged ipsilateral to the TMS site. This provides further evidence for a link between attention and intrinsic theta rhythmicity in a fronto-parietal attention network.

Fiebelkorn and Kastner’s rhythmic theory of attention (Fiebelkorn & Kastner, 2020) posits that optimal theta phase associated with higher performance and attentional focus is linked to enhanced gamma power in occipito-parietal areas. Although we did not find modulation of gamma power, we found a modulation of gamma band variability. Spatial attention was associated with less gamma band variability in the stimulated left IPS and, potentially, in posterior occipital regions when attention was directed contralaterally rather than ipsilaterally to the probe pulse. Increased power and phase consistency of gamma oscillations have repeatedly been shown in sensory areas during manipulations of attention allocation (Bichot et al., 2005; Fries, 2009). These gamma modulations are thought to reflect increased feed-forward processing of sensory stimuli (Bastos et al., 2015; van Kerkoerle et al., 2014). Here, we observed decreased gamma band phase variability even in the absence of a novel visual stimulus to be processed, and while peripheral stimuli were identical across hemifields, indicating that the decreased phase variability may be the result of changes in local IPS network properties rather than the propagation of visual evoked responses occurring at lower levels of the cortical hierarchy. These results suggest that left parieto-occipital areas have a greater propensity for gamma oscillation reset when spatial attention is engaged contralaterally rather than ipsilaterally. This is interesting because it provides complementary evidence that intrinsic network properties are modulated during attentional allocation (Fries, 2009), not only in visual sensory areas but also in higher parietal cortex.

In this study, we stimulated the left IPS because it has been shown to be implicated in the maintenance of spatial attention and to display strong contralaterally-biased activity (Jeong & Xu, 2016; Sheremata et al., 2010, 2018; Szczepanski et al., 2010). Unlike Blankenburg and colleagues (Blankenburg et al., 2010), who reported a bilateral influence of right parietal TMS on visual areas during the spatial deployment of attention, we did not observe bilateral changes in variability (ITPC) following left IPS stimulation. Instead, we observed reduced gamma and theta band phase variability only in left occipito-parietal areas when attention was allocated to the right side of space (contralateral to the side of TMS). This finding is compatible with the hypothesis that while the right IPS controls attentional deployment over both the left and right sides of space, the left IPS has a more restricted role, controlling only attentional allocation over the right side of space (Capotosto et al., 2012). Likewise, the right FEF is suggested to be dominant over the left in controlling the allocation of visuo-spatial attention (Murd et al., 2020). Here, we observed reduced theta variability over the right frontal region. From this result alone, we cannot conclude whether the frontal cluster forms part of a network with contralateral IPS, or whether the right (rather than left) frontal region is engaged with both parietal regions. This ambiguity could be addressed by stimulating the right IPS under the same experimental conditions as those used in the current study. If the network is contralateralised, then decreased theta variability and increased TEP amplitude should be seen in the left frontal region when participants attend to the left side of space, relative to the right side of space. If the right frontal region is dominant in its connection with the posterior IPS, then the same decreased theta variability and increased TEP amplitude found in the current study should be seen in right frontal areas when participants attend to the left side of space, relative to the right side of space.

### Attentional state can be decoded following stimulation of both IPS and S1

Here, we sought to characterise participants’ attentional state, as defined by the location of spatial attention and perceptual load, using multivariate decoding of TMS-evoked potentials (Rose et al., 2016). The ability to decode attention states in response to the TMS probe indicated that stimulation uncovered information about the experimental condition stored in whole-scalp EEG data that was not decodable in the baseline period. Given the fact that task difficulty modulated TEP amplitude after both IPS and S1 stimulation (Fig.3), it is not surprising that task difficulty was decodable from TEP amplitude after both IPS and S1 TMS. Perhaps more surprising was the ability to decode attention side after both IPS and S1 TMS, even though no change in TEP amplitude was detected in the univariate analyses for S1. This finding suggests that uncovering latent information following a TMS pulse is not specific to stimulation of the network engaged in the task. A lack of stimulation-site specificity is consistent with previous findings. Rose and colleagues (Rose et al., 2016) used multivariate decoding of EEG data to determine the category (word, face or motion signal) of two items held in working memory. The items could be either attended or unattended in working memory, but both were relevant at some point in the trial. Results showed that the category of unattended items stored in working memory could be recovered from neural activity following a TMS pulse. Interestingly, the TMS pulse did not have to be applied to the category-specific brain region to uncover the information. How might this happen? One possibility is that stimulation of S1 recruited a distant cortical area that was also recruited by IPS stimulation, and whose activity was actually modulated by the task. For example, both S1 (Rolls et al., 2023) and IPS (Alahmadi, 2021) have connections to areas BA5 and BA7 in the parietal cortex. In this case, our decoding result may indicate that TMS probes heightened connectivity across distributed networks that share key nodes representing task-relevant activity. Another possibility relates to the known interplay between attention and sensory signals. The sensory aspects of a TMS pulse (auditory and somatosensory) result in peripheral evoked potentials (Conde et al., 2019; Rocchi et al., 2021; Siebner et al., 2019), which could have been modulated by attentional state and the classifier could have used this non-specific information to classify trials. While this seems possible for the attentional load conditions, it does not easily account for the variable timing of decoding effects across S1 and IPS (Fig.6). Intriguingly, non-specific sensory input (e.g., a high contrast visual stimulus) has also been shown to boost the decodability of items held in working memory (Wolff et al., 2017), albeit likely through a different mechanism to TMS-evoked decoding boosts (Barbosa et al., 2021). It is therefore possible that a TMS pulse can reveal hidden state information through several pathways, some network-specific and some non-specific.

### Task difficulty increases the amplitude of site-non-specific potentials

The amplitude of a large vertex positive potential around 180ms was significantly larger in the Hard than in the Easy visual-motion condition. This was true for both the peak analysis and the whole scalp analysis (Fig.3). Notably, this potential occurred after both IPS and S1 TMS and was modulated in a similar way for both TMS sites. While it is possible that stimulation of S1 led to activity in a distant brain region that is also engaged by IPS TMS, it seems more likely that the P181 potential is a multimodal response to the audio-somatosensory aspects of TMS. Indeed, previous work investigating the multisensory origin of TEPs has highlighted the N1-P2 complex (N100-P180) as being a result of auditory and somatosensory stimulation, inviting caution in its interpretation (Conde et al., 2019; Rocchi et al., 2021; Siebner et al., 2019). While we followed best practice by masking the TMS coil ‘click’ with in-ear white noise (Hernandez-Pavon et al., 2023; Rocchi et al., 2021), we could only deliver the white noise at a level comfortable for participants, which was likely not sufficient to fully mask TMS noise given the stimulation intensity (100% of RMT). The resulting vertex ‘alerting’ potential has been linked to the processing of salient stimuli in several modalities (Mouraux & Iannetti, 2009). Why was this potential increased in the difficult (high-load) condition? In our study, any TMS sensory stimuli could be considered distractors relative to the main visual attention task. The cross-modal attention literature indicates that irrelevant distractor processing is often inhibited when participants focus on a primary task (Brockhoff et al., 2022). However, increased distractor processing has also been reported when the main task draws too much on cognitive control resources (Regenbogen et al., 2012), which may have been the case in the high-load condition of our study. Methodological advances using machine learning (Cristofari et al., 2023) may one day allow us to remove multimodal responses from TMS-EEG data. Until these techniques generalize to newly acquired data, however, our results highlight the importance of a realistic or active sham control that can match artefacts in the experimental condition of interest.

## Conclusions

Here we have shown that TMS-evoked responses originating from the left IPS are modulated under different attentional states. Amplitude modulations of TEPs were often minimal and non-specific to the stimulation site. In contrast, the variability of theta and gamma oscillatory phase was reduced during spatial attention maintenance in a parieto-frontal network, confirming the importance of these oscillatory processes in attention and providing evidence for a dynamic adjustment of IPS network properties according to attentional state.

## Data and code availability statement

De-identified data is available upon reasonable request; code used for stimulus presentation, data collection, and data analysis is available at the Open Science Framework (https://osf.io/axnyk/).

## Acknowledgements

We thank Dr. David Lloyd for technical assistance, Miss Imogen Stead and Miss Sylvie Loneragan for data acquisition and Dr. Hamid Karimi Rouzbahani and all Mattingley lab members for helpful discussions.

## Notes

**Conflicts of interest:** None to declare

**Funding:** This work was supported by a grant from the National Health and Medical Research Council (NHMRC) Australia (GNT1129715) awarded to JBM and PED. JBM was supported by an NHMRC Investigator Grant (GNT2010141).

### Competing Interest Statement

The authors have declared no competing interest.

## References

Alahmadi, A. A. S. (2021). Investigating the sub-regions of the superior parietal cortex using functional magnetic resonance imaging connectivity. Insights into Imaging, 12(1), 47. 10.1186/s13244-021-00993-9

Arazi, A., Yeshurun, Y., & Dinstein, I. (2019). Neural Variability Is Quenched by Attention. The Journal of Neuroscience: The Official Journal of the Society for Neuroscience, 39(30), 5975–5985. 10.1523/JNEUROSCI.0355-19.2019

Barbosa, J., Lozano-Soldevilla, D., & Compte, A. (2021). Pinging the brain with visual impulses reveals electrically active, not activity-silent, working memories. PLoS Biology, 19(10), e3001436. 10.1371/journal.pbio.3001436

Bastos, A. M., Vezoli, J., Bosman, C. A., Schoffelen, J.-M., Oostenveld, R., Dowdall, J. R., De Weerd, P., Kennedy, H., & Fries, P. (2015). Visual Areas Exert Feedforward and Feedback Influences through Distinct Frequency Channels. Neuron, 85(2), 390–401. 10.1016/j.neuron.2014.12.018

Bergmann, T. O., Karabanov, A., Hartwigsen, G., Thielscher, A., & Siebner, H. R. (2016). Combining non-invasive transcranial brain stimulation with neuroimaging and electrophysiology: Current approaches and future perspectives. NeuroImage, 140, 4–19. 10.1016/j.neuroimage.2016.02.012

Bichot, N. P., Rossi, A. F., & Desimone, R. (2005). Parallel and Serial Neural Mechanisms for Visual Search in Macaque Area V4. Science, 308(5721), 529–534. 10.1126/science.1109676

Blankenburg, F., Ruff, C. C., Bestmann, S., Bjoertomt, O., Josephs, O., Deichmann, R., & Driver, J. (2010). Studying the Role of Human Parietal Cortex in Visuospatial Attention with Concurrent TMS–fMRI. Cerebral Cortex, 20(11), 2702–2711. 10.1093/cercor/bhq015

Bradley, C., Nydam, A. S., Dux, P. E., & Mattingley, J. B. (2022). State-dependent effects of neural stimulation on brain function and cognition. Nature Reviews Neuroscience.

Brockhoff, L., Schindler, S., Bruchmann, M., & Straube, T. (2022). Effects of perceptual and working memory load on brain responses to task-irrelevant stimuli: Review and implications for future research. Neuroscience & Biobehavioral Reviews, 135, 104580. 10.1016/j.neubiorev.2022.104580

Capotosto, P., Babiloni, C., Romani, G. L., & Corbetta, M. (2009). Frontoparietal Cortex Controls Spatial Attention through Modulation of Anticipatory Alpha Rhythms. Journal of Neuroscience, 29(18), 5863–5872. 10.1523/JNEUROSCI.0539-09.2009

Capotosto, P., Babiloni, C., Romani, G. L., & Corbetta, M. (2012). Differential Contribution of Right and Left Parietal Cortex to the Control of Spatial Attention: A Simultaneous EEG-rTMS Study. Cerebral Cortex, 22(2), 446–454. 10.1093/cercor/bhr127

Capotosto, P., Spadone, S., Tosoni, A., Sestieri, C., Romani, G. L., Della Penna, S., & Corbetta, M. (2015). Dynamics of EEG Rhythms Support Distinct Visual Selection Mechanisms in Parietal Cortex: A Simultaneous Transcranial Magnetic Stimulation and EEG Study. The Journal of Neuroscience, 35(2), 721–730. 10.1523/JNEUROSCI.2066-14.2015

Cohen. (2014). Analyzing Neural Time Series Data. MIT Press. https://mitpress.mit.edu/9780262019873/analyzing-neural-time-series-data/

Conde, V., Tomasevic, L., Akopian, I., Stanek, K., Saturnino, G. B., Thielscher, A., Bergmann, T. O., & Siebner, H. R. (2019). The non-transcranial TMS-evoked potential is an inherent source of ambiguity in TMS-EEG studies. NeuroImage, 185, 300–312. 10.1016/j.neuroimage.2018.10.052

Corbetta, M., Patel, G., & Shulman, G. L. (2008). The Reorienting System of the Human Brain: From Environment to Theory of Mind. Neuron, 58(3), 306–324. 10.1016/j.neuron.2008.04.017

Corbetta, M., & Shulman, G. L. (2002). Control of goal-directed and stimulus-driven attention in the brain. Nature Reviews Neuroscience, 3(3), Article 3. 10.1038/nrn755

Cristofari, A., De Santis, M., Lucidi, S., Rothwell, J., Casula, E. P., & Rocchi, L. (2023). Machine Learning-Based Classification to Disentangle EEG Responses to TMS and Auditory Input. Brain Sciences, 13(6), 866. 10.3390/brainsci13060866

Fiebelkorn, I. C., & Kastner, S. (2020). Functional Specialization in the Attention Network. Annual Review of Psychology, 71(1), 221–249. 10.1146/annurev-psych-010418-103429

Fries, P. (2009). Neuronal Gamma-Band Synchronization as a Fundamental Process in Cortical Computation. Annual Review of Neuroscience, 32(1), 209–224. 10.1146/annurev.neuro.051508.135603

Gramfort, A., Luessi, M., Larson, E., Engemann, D., Strohmeier, D., Brodbeck, C., Goj, R., Jas, M., Brooks, T., Parkkonen, L., & Hämäläinen, M. (2013). MEG and EEG data analysis with MNE-Python. Frontiers in Neuroscience, 7. https://www.frontiersin.org/articles/10.3389/fnins.2013.00267

Grootswagers, T., Wardle, S. G., & Carlson, T. A. (2017). Decoding Dynamic Brain Patterns from Evoked Responses: A Tutorial on Multivariate Pattern Analysis Applied to Time Series Neuroimaging Data. Journal of Cognitive Neuroscience, 29(4), 677–697. 10.1162/jocn_a_01068

Heinen, K., Feredoes, E., Weiskopf, N., Ruff, C. C., & Driver, J. (2014). Direct evidence for attention-dependent influences of the frontal eye-fields on feature-responsive visual cortex. Cerebral Cortex (New York, N.Y.: 1991), 24(11), 2815–2821. 10.1093/cercor/bht157

Hernandez-Pavon, J. C., Veniero, D., Bergmann, T. O., Belardinelli, P., Bortoletto, M., Casarotto, S., Casula, E. P., Farzan, F., Fecchio, M., Julkunen, P., Kallioniemi, E., Lioumis, P., Metsomaa, J., Miniussi, C., Mutanen, T. P., Rocchi, L., Rogasch, N. C., Shafi, M. M., Siebner, H. R., … Ilmoniemi, R. J. (2023). TMS combined with EEG: Recommendations and open issues for data collection and analysis. Brain Stimulation, 16(2), 567–593. 10.1016/j.brs.2023.02.009

Herring, J. D., Thut, G., Jensen, O., & Bergmann, T. O. (2015). Attention Modulates TMS-Locked Alpha Oscillations in the Visual Cortex. Journal of Neuroscience, 35(43), 14435–14447. 10.1523/JNEUROSCI.1833-15.2015

Hussain, S. J., & Quentin, R. (2022). Decoding personalized motor cortical excitability states from human electroencephalography. Scientific Reports, 12(1), Article 1. 10.1038/s41598-022-10239-3

Jeong, S. K., & Xu, Y. (2016). The impact of top-down spatial attention on laterality and hemispheric asymmetry in the human parietal cortex. Journal of Vision. 10.1167/16.10.2

Keel, J. C., Smith, M. J., & Wassermann, E. M. (2001). A safety screening questionnaire for transcranial magnetic stimulation. Clinical Neurophysiology, 112(4), 720. 10.1016/S1388-2457(00)00518-6

Kelley, T. A., Serences, J. T., Giesbrecht, B., & Yantis, S. (2008). Cortical Mechanisms for Shifting and Holding Visuospatial Attention. Cerebral Cortex, 18(1), 114–125. 10.1093/cercor/bhm036

Larson, E., Gramfort, A., Engemann, D. A., Leppakangas, J., Brodbeck, C., Jas, M., Brooks, T., Sassenhagen, J., Luessi, M., McCloy, D., King, J.-R., Höchenberger, R., Goj, R., Favelier, G., Brunner, C., van Vliet, M., Wronkiewicz, M., Holdgraf, C., Rockhill, A., … luzpaz. (2024). MNE-Python (v1.6.1) [Computer software]. Zenodo. 10.5281/zenodo.10519948

Lee, M. D., & Wagenmakers, E.-J. (2014). Bayesian Cognitive Modeling: A Practical Course. Cambridge University Press. 10.1017/CBO9781139087759

Leitão, J., Thielscher, A., Tünnerhoff, J., & Noppeney, U. (2015). Concurrent TMS-fMRI Reveals Interactions between Dorsal and Ventral Attentional Systems. The Journal of Neuroscience: The Official Journal of the Society for Neuroscience, 35(32), 11445– 11457. 10.1523/JNEUROSCI.0939-15.2015

Maris, E., & Oostenveld, R. (2007). Nonparametric statistical testing of EEG- and MEG-data. Journal of Neuroscience Methods, 164(1), 177–190. 10.1016/j.jneumeth.2007.03.024

Marshall, T. R., O’Shea, J., Jensen, O., & Bergmann, T. O. (2015). Frontal Eye Fields Control Attentional Modulation of Alpha and Gamma Oscillations in Contralateral Occipitoparietal Cortex. Journal of Neuroscience, 35(4), 1638–1647. 10.1523/JNEUROSCI.3116-14.2015

Metsomaa, J., Belardinelli, P., Ermolova, M., Ziemann, U., & Zrenner, C. (2021). Causal decoding of individual cortical excitability states. NeuroImage, 245, 118652. 10.1016/j.neuroimage.2021.118652

Michel, C. M., & He, B. (2019). EEG source localization. Handbook of Clinical Neurology, 160, 85–101. 10.1016/B978-0-444-64032-1.00006-0

Morey, R. D., & Rouder, J. N. (2018). Baysefactor: Computation of Bayes Factors for Common Designs.

Morishima, Y., Akaishi, R., Yamada, Y., Okuda, J., Toma, K., & Sakai, K. (2009). Task-specific signal transmission from prefrontal cortex in visual selective attention. Nature Neuroscience, 12(1), Article 1. 10.1038/nn.2237

Morris, A. P., Chambers, C. D., & Mattingley, J. B. (2007). Parietal stimulation destabilizes spatial updating across saccadic eye movements. Proceedings of the National Academy of Sciences, 104(21), 9069–9074. 10.1073/pnas.0610508104

Mouraux, A., & Iannetti, G. D. (2009). Nociceptive Laser-Evoked Brain Potentials Do Not Reflect Nociceptive-Specific Neural Activity. Journal of Neurophysiology, 101(6), 3258–3269. 10.1152/jn.91181.2008

Murd, C., Moisa, M., Grueschow, M., Polania, R., & Ruff, C. C. (2020). Causal contributions of human frontal eye fields to distinct aspects of decision formation. Scientific Reports, 10. 10.1038/s41598-020-64064-7

Noreika, V., Kamke, M. R., Canales-Johnson, A., Chennu, S., Bekinschtein, T. A., & Mattingley, J. B. (2020). Alertness fluctuations when performing a task modulate cortical evoked responses to transcranial magnetic stimulation. NeuroImage, 223, 117305. 10.1016/j.neuroimage.2020.117305

Okazaki, Y. O., Mizuno, Y., & Kitajo, K. (2020). Probing dynamical cortical gating of attention with concurrent TMS-EEG. Scientific Reports, 10(1), 4959. 10.1038/s41598-020-61590-2

Oostenveld, R., Fries, P., Maris, E., & Schoffelen, J.-M. (2011). FieldTrip: Open Source Software for Advanced Analysis of MEG, EEG, and Invasive Electrophysiological Data. Computational Intelligence and Neuroscience, 2011, 1–9. 10.1155/2011/156869

Oosterhof, N. N., Connolly, A. C., & Haxby, J. V. (2016). CoSMoMVPA: Multi-Modal Multivariate Pattern Analysis of Neuroimaging Data in Matlab/GNU Octave. Frontiers in Neuroinformatics, 10, 27. 10.3389/fninf.2016.00027

Ozdemir, R. A., Tadayon, E., Boucher, P., Momi, D., Karakhanyan, K. A., Fox, M. D., Halko, M. A., Pascual-Leone, A., Shafi, M. M., & Santarnecchi, E. (2020). Individualized perturbation of the human connectome reveals reproducible biomarkers of network dynamics relevant to cognition. Proceedings of the National Academy of Sciences, 117(14), 8115–8125. 10.1073/pnas.1911240117

Ozdemir, R. A., Tadayon, E., Boucher, P., Sun, H., Momi, D., Ganglberger, W., Westover, M. B., Pascual-Leone, A., Santarnecchi, E., & Shafi, M. M. (2021). Cortical responses to noninvasive perturbations enable individual brain fingerprinting. Brain Stimulation, 14(2), 391–403. 10.1016/j.brs.2021.02.005

Pashler, H. E. (1998). The psychology of attention (pp. xiv, 494). The MIT Press.

Poghosyan, V., & Ioannides, A. A. (2008). Attention modulates earliest responses in the primary auditory and visual cortices. Neuron, 58(5), 802–813. 10.1016/j.neuron.2008.04.013

Polanía, R., Nitsche, M. A., & Ruff, C. C. (2018). Studying and modifying brain function with non-invasive brain stimulation. Nature Neuroscience, 21(2), Article 2. 10.1038/s41593-017-0054-4

Regenbogen, C., Vos, M. D., Debener, S., Turetsky, B. I., Mößnang, C., Finkelmeyer, A., Habel, U., Neuner, I., & Kellermann, T. (2012). Auditory Processing under Cross-Modal Visual Load Investigated with Simultaneous EEG-fMRI. PLOS ONE, 7(12), e52267. 10.1371/journal.pone.0052267

Riddle, J., Hwang, K., Cellier, D., Dhanani, S., & D’Esposito, M. (2019). Causal Evidence for the Role of Neuronal Oscillations in Top–Down and Bottom–Up Attention. Journal of Cognitive Neuroscience, 31(5), 768–779. 10.1162/jocn_a_01376

Rocchi, L., Santo, A. D., Brown, K., Ibáñez, J., Casula, E., Rawji, V., Lazzaro, V. D., Koch, G., & Rothwell, J. (2021). Disentangling EEG responses to TMS due to cortical and peripheral activations. *Brain Stimulation: Basic*, Translational, and Clinical Research in Neuromodulation, 14(1), 4–18. 10.1016/j.brs.2020.10.011

Rogasch, N. C., Sullivan, C., Thomson, R. H., Rose, N. S., Bailey, N. W., Fitzgerald, P. B., Farzan, F., & Hernandez-Pavon, J. C. (2017). Analysing concurrent transcranial magnetic stimulation and electroencephalographic data: A review and introduction to the open-source TESA software. NeuroImage, 147, 934–951. 10.1016/j.neuroimage.2016.10.031

Rolls, E. T., Deco, G., Huang, C.-C., & Feng, J. (2023). Prefrontal and somatosensory-motor cortex effective connectivity in humans. Cerebral Cortex, 33(8), 4939–4963. 10.1093/cercor/bhac391

Romei, V., Thut, G., Mok, R. M., Schyns, P. G., & Driver, J. (2012). Causal implication by rhythmic transcranial magnetic stimulation of alpha frequency in feature-based local vs. Global attention. The European Journal of Neuroscience, 35(6), 968–974. 10.1111/j.1460-9568.2012.08020.x

Rosanova, M., Casali, A., Bellina, V., Resta, F., Mariotti, M., & Massimini, M. (2009). Natural Frequencies of Human Corticothalamic Circuits. Journal of Neuroscience, 29(24), 7679–7685. 10.1523/JNEUROSCI.0445-09.2009

Rose, N. S., LaRocque, J. J., Riggall, A. C., Gosseries, O., Starrett, M. J., Meyering, E. E., & Postle, B. R. (2016). Reactivation of latent working memories with transcranial magnetic stimulation. Science, 354(6316), 1136–1139. 10.1126/science.aah7011

Rossini, P. M., Burke, D., Chen, R., Cohen, L. G., Daskalakis, Z., Di Iorio, R., Di Lazzaro, V., Ferreri, F., Fitzgerald, P. B., George, M. S., Hallett, M., Lefaucheur, J. P., Langguth, B., Matsumoto, H., Miniussi, C., Nitsche, M. A., Pascual-Leone, A., Paulus, W., Rossi, S., … Ziemann, U. (2015). Non-invasive electrical and magnetic stimulation of the brain, spinal cord, roots and peripheral nerves: Basic principles and procedures for routine clinical and research application. An updated report from an I.F.C.N. Committee. Clinical Neurophysiology, 126(6), 1071–1107. 10.1016/j.clinph.2015.02.001

Sassenhagen, J., & Draschkow, D. (2019). Cluster-based permutation tests of MEG/EEG data do not establish significance of effect latency or location. Psychophysiology, 56(6), e13335. 10.1111/psyp.13335

Sheremata, S. L., Bettencourt, K. C., & Somers, D. C. (2010). Hemispheric asymmetry in visuotopic posterior parietal cortex emerges with visual short-term memory load. The Journal of Neuroscience: The Official Journal of the Society for Neuroscience, 30(38), 12581–12588. 10.1523/JNEUROSCI.2689-10.2010

Sheremata, S. L., Somers, D. C., & Shomstein, S. (2018). Visual Short-Term Memory Activity in Parietal Lobe Reflects Cognitive Processes beyond Attentional Selection. The Journal of Neuroscience: The Official Journal of the Society for Neuroscience, 38(6), 1511–1519. 10.1523/JNEUROSCI.1716-17.2017

Siebner, H. R., Conde, V., Tomasevic, L., Thielscher, A., & Bergmann, T. O. (2019). Distilling the essence of TMS-evoked EEG potentials (TEPs): A call for securing mechanistic specificity and experimental rigor. Brain Stimulation: Basic, Translational, and Clinical Research in Neuromodulation, 12(4), 1051–1054. 10.1016/j.brs.2019.03.076

Silver, M. A., & Kastner, S. (2009). Topographic maps in human frontal and parietal cortex. Trends in Cognitive Sciences, 13(11), 488–495. 10.1016/j.tics.2009.08.005

Skrandies, W. (1990). Global field power and topographic similarity. Brain Topography, 3(1), 137–141. 10.1007/BF01128870

Speed, A., Del Rosario, J., Mikail, N., & Haider, B. (2020). Spatial attention enhances network, cellular and subthreshold responses in mouse visual cortex. Nature Communications, 11(1), 505. 10.1038/s41467-020-14355-4

Stokes, M. G., Chambers, C. D., Gould, I. C., Henderson, T. R., Janko, N. E., Allen, N. B., & Mattingley, J. B. (2005). Simple Metric For Scaling Motor Threshold Based on Scalp-Cortex Distance: Application to Studies Using Transcranial Magnetic Stimulation. Journal of Neurophysiology, 94(6), 4520–4527. 10.1152/jn.00067.2005

Szczepanski, S. M., Konen, C. S., & Kastner, S. (2010). Mechanisms of Spatial Attention Control in Frontal and Parietal Cortex. Journal of Neuroscience, 30(1), 148–160. 10.1523/JNEUROSCI.3862-09.2010

Teichmann, L., Moerel, D., Baker, C., & Grootswagers, T. (2022). An Empirically Driven Guide on Using Bayes Factors for M/EEG Decoding. Aperture Neuro, 2022(8), 52. 10.52294/82179f90-eeb9-4933-adbe-c2a454577289

Tosoni, A., Capotosto, P., Baldassarre, A., Spadone, S., & Sestieri, C. (2023). Neuroimaging evidence supporting a dual-network architecture for the control of visuospatial attention in the human brain: A mini review. Frontiers in Human Neuroscience, 17. https://www.frontiersin.org/articles/10.3389/fnhum.2023.1250096

van Diepen, R. M., & Mazaheri, A. (2018). The Caveats of observing Inter-Trial Phase-Coherence in Cognitive Neuroscience. Scientific Reports, 8(1), 2990. 10.1038/s41598-018-20423-z

van Kerkoerle, T., Self, M. W., Dagnino, B., Gariel-Mathis, M.-A., Poort, J., van der Togt, C., & Roelfsema, P. R. (2014). Alpha and gamma oscillations characterize feedback and feedforward processing in monkey visual cortex. Proceedings of the National Academy of Sciences, 111(40), 14332–14341. 10.1073/pnas.1402773111

Wang, M., Yu, B., Luo, C., Fogelson, N., Zhang, J., Jin, Z., & Li, L. (2020). Evaluating the causal contribution of fronto-parietal cortices to the control of the bottom-up and top-down visual attention using fMRI-guided TMS. Cortex, 126, 200–212. 10.1016/j.cortex.2020.01.005

Wolff, M. J., Jochim, J., Akyürek, E. G., & Stokes, M. G. (2017). Dynamic hidden states underlying working-memory-guided behavior. Nature Neuroscience, 20(6), Article 6. 10.1038/nn.4546

Xu, J., Calhoun, V. D., Pearlson, G. D., & Potenza, M. N. (2014). Opposite Modulation of Brain Functional Networks Implicated at Low vs. High Demand of Attention and Working Memory. PLOS ONE, 9(1), e87078. 10.1371/journal.pone.0087078

